# An optimized live bacterial delivery platform for the production and delivery of therapeutic nucleic acids and proteins

**DOI:** 10.1101/2021.10.17.464697

**Authors:** Darcy S.O. Mora, Madeline Cox, Forgivemore Magunda, Ashley B. Williams, Lyndsey Linke

## Abstract

There is an unmet need for delivery platforms that realize the full potential of next-generation therapeutic and vaccine technologies, especially those that require intracellular delivery of nucleic acids. The in vivo usefulness of the current state-of-the-art delivery systems is limited by numerous intrinsic weaknesses, including lack of targeting specificity, inefficient entry and endosomal escape into target cells, undesirable immune activation, off-target effects, a small therapeutic window, limited genetic encoding and cargo capacity, and manufacturing challenges. Here we present our characterization of a delivery platform based on the use of engineered live, tissue-targeting, non-pathogenic bacteria (*Escherichia coli* strain SVC1) for intracellular cargo delivery. The SVC1 bacteria are engineered to specifically bind to epithelial cells via a surface-expressed targeting ligand, to escape the endosome upon intracellularization, and to have minimal immunogenicity. Here we report findings on key features of this system. First, we demonstrated that bacterial delivery of a short hairpin RNA (shRNA) can target and silence a gene in an in vitro mammalian respiratory cell model. Next, we used an in vivo mouse model to demonstrate that SVC1 bacteria are invasive to epithelial cells of various tissues and organs (eye, nose, mouth, stomach, vagina, skeletal muscle, and lungs) via local administration. We also showed that repeat dosing of SVC1 bacteria to the lungs is minimally immunogenic and that it does not have adverse effects on tissue homeostasis. Finally, to validate the potential of SVC1 bacteria in therapeutic applications, we demonstrated that bacterial delivery of influenza-targeting shRNAs to the respiratory tissues can mitigate viral replication in a mouse model of influenza infection. Our ongoing work is focused on further refining this platform for efficient delivery of nucleic acids, gene editing machinery, and therapeutic proteins, and we expect that this platform technology will enable a wide range of advanced therapeutic approaches.

## Introduction

Nucleic acid and protein moieties have rapidly come to the forefront of biomedical research as alternatives to small molecules and other drugs for the treatment of a variety of human diseases as well as a powerful approach for developing next-generation vaccines. To fully realize the potential of these therapeutic and vaccination approaches, proteins and nucleic acids must be delivered to target cells. Currently, such delivery in patients is mostly limited to viral vectors and other chemical delivery modalities (e.g., lipid nanoparticles); however, the power of these approaches is limited by a number of drawbacks, including safety concerns, production cost, payload size limitations, immunogenicity, and cytotoxicity ^1–5^. To overcome these limitations, alternative delivery systems must be developed. For over three decades, there has been increasing interest in the use of engineered commensal or probiotic bacteria as an in vivo nucleic acid and protein delivery modality as they represent a powerful and unique approach for the delivery of therapeutic moieties that can overcome many of the limitations of viral or chemical delivery systems. As part of the normal human flora, *Escherichia coli* has garnered particular interest ^6–13^.

To serve as a viable in vivo delivery approach, a bacteria*-*based delivery system must possess three key activities: 1) the bacterial cells must be able to specifically target a disease-relevant cell type (e.g., epithelial cells); 2) the bacteria must be able to enter the target cell upon arrival; and 3) the bacteria must be able to escape the endosome to deposit their cargo into the cytoplasm. While professional phagocytic cells (e.g., neutrophils) readily take up bacteria via phagocytosis ^14^, other cell types, which are clear front runners as therapeutic targets (e.g., epithelial cells), do not have intrinsic phagocytic activity; thus, the bacteria must be engineered to enable cellular uptake. In the *Escherichia coli*-based system described here, the *E. coli* strain SVC1 has been genetically modified to meet these needs ^9^. First, the SVC1 bacteria carry a heterologous gene encoding the *Yersinia pseudotuberculosis* invasin (inv) protein to allow uptake by the targeted eukaryotic cells. Invasin binds to β1 integrin ^15–18^, which occurs with several integrin receptors (α3, α4, α5, α6, and αv) ^19^. The specificity for β1 integrin allows the bacteria to target only β1-expressing cells ^15^. Important for the usefulness of an invasin-based targeting system is that β1 integrin is expressed by a wide variety of cells, including a broad group of epithelial cells, e.g., intestinal ^20^, respiratory ^21^, eye ^22^, and others ^19^, as a well as certain types of cancer cells ^23,24^. Through local or systemic administration, the bacterial delivery vehicle can target various organs and tissues after which it interacts with specific types of epithelial or cancerous cells. Upon binding to β1 integrin on the target cell surface, the bacteria are taken up via receptor-mediated endocytosis, thereafter finding themselves compartmentalized within endosomes.

For the cargo to reach the cytoplasm of the target cell, the cargo must be released from both the bacterial cell and the target cell’s endosome. These steps are enabled by two additional modifications to the SVC1 bacteria. First, the bacteria are auxotrophic for diaminopimelic acid (DAP) due to a null mutation in *dapA*, which encodes a protein essential for DAP biosynthesis; thus, DAP must be provided exogenously in the growth medium. As DAP is required for peptidoglycan crosslinking in the Gram-negative cell wall, bacteria are structurally compromised in the absence of DAP ^13,25^. Eukaryotic cells do not produce DAP, so upon entry the bacterial cells 1) cannot replicate since they cannot synthesize new cell wall and 2) are sufficiently unstable that they rapidly lyse to release their cargo into the endosome. Upon lysis in the endosome, the bacteria release the product of a second heterologous gene, *hlyA*. This gene encodes the hemolysin HlyA from *Listeria monocytogenes* (also known as listeriolysin O or LLO) ^26^. The liberated LLO protein perforates the endosomal membrane, thereby allowing the bacterial cargo to enter the cytoplasm of the eukaryotic cell ^13,27,28^. The combined traits conferred by these genetic modifications render these bacteria useful as an efficient intracellular delivery vehicle.

An important consideration of any drug delivery vehicle that will be used in therapeutic applications is safety. Of particular relevance to bacterial drug delivery vehicles are the safety aspects of host colonization and immunogenicity. The parental strain of SVC1 has been highly domesticated since its isolation from a human patient in 1922 ^29^. The lipopolysaccharide core on the outer membrane is defective in attachment of the O-antigen, making SVC1 a rough strain. Furthermore, the cells do not express a capsular (K) antigen, further inhibiting colonization ^25^. As mentioned above, the bacteria described here are attenuated via engineered DAP auxotrophy that, in addition to promoting cargo release in the endosome, also acts as a powerful biocontainment mechanism, both in the patient and in the environment ^25^. To support bacterial growth and replication, supplemental DAP must be provided in the bacterial growth medium. As DAP is absent from the cytoplasm of eukaryotic cells, these SVC1 bacteria cannot replicate within the target cell. Furthermore, outside of a very low level of DAP in the urine (likely originating from lysed bacteria in the excretory system) ^30^, DAP is not otherwise known to exist in the human body or inside cells. Thus, SVC1 bacteria cannot replicate in patient tissues or within cells. Another additional safety concern is horizontal gene transfer (HGT), whereby genes that confer increased fitness under environmental conditions (e.g., antibiotic resistance genes) are passed from a donor strain to a compatible recipient strain ^31^. HGT is mediated by a variety of plasmids and phages; importantly, the strain described here lacks such plasmids and phage ^32^, preventing it from disseminating cargo-encoding recombinant molecules.

Current delivery systems, including lipid nanoparticles and viral vectors, suffer from a variety of immunological issues ^33–40^ and the outcomes of undesirable immune responses can have devastating clinical repercussions, especially in applications requiring repeat administration. Therefore, the immunogenicity of any novel delivery system must be carefully characterized to mitigate such issues. As part of the normal flora, *E. coli* represents an appealing candidate for use as a bacteria-based delivery vehicle, particularly in light of immunogenicity concerns; nevertheless, the immunogenicity of a bacterial system must be empirically examined in vivo.

As highlighted by the recent surge in the use of mRNA-based vaccines to combat the SARS-CoV-2 pandemic, the intracellular delivery of nucleic acids (RNA in particular) has garnered significant attention due to its remarkable therapeutic potential ^41,42^. While the most clinically advanced application of nucleic acid delivery currently in use is indeed for vaccination, there has been an ongoing interest in the use of RNA interference (RNAi) as a therapeutic modality in other clinical applications ^43–45^. The mechanism underlying RNAi therapy depends on intracellular delivery of an RNA molecule to a target cell that can then be processed by the cellular RNAi pathway to ultimately silence the expression of a specifically targeted pathological gene. RNAi-mediated silencing is guided by small interfering RNAs (siRNAs)^46–48^. Substrate siRNAs can be generated from the enzymatic processing of short hairpin RNAs (shRNAs), which, importantly, can be expressed by bacteria ^9,28^. *E. coli* strains can be engineered to express shRNAs against a target gene of interest. In the case of invasive strains, e.g., SVC1, the bacteria can both produce and deliver the shRNAs to the cytoplasm of targeted cells to achieve desired therapeutic effects. Some progress has been made in the therapeutic use of shRNA-expressing *E. coli* strains. For example, a therapeutic *E. coli* strain was generated as a treatment for familial adenomatous polypopsis (FAP), a disease of the gastrointestinal tract, via silencing of endogenous beta-catenin in gastrointestinal epithelial cells ^28^. Related preclinical and human clinical studies demonstrated the safety and efficacy of this approach, highlighting the potential of *E. coli-*mediated nucleic acid delivery for development in other clinical applications ^10,11,49^. Linke and colleagues developed an shRNA-based approach for the prophylaxis and treatment of avian influenza virus ^9^. In this system, two *E. coli* strains were engineered to independently silence the expression of two essential influenza viral proteins: an RNA polymerase subunit (polymerase acidic protein; PA) and a capsid protein (nucleoprotein; NP). When administered together to the respiratory tract, these bacterial strains can target most known influenza A strains while minimizing the development of therapeutic resistance, especially that resulting from genetic drift ^50–53^. The safety and efficacy of this approach were robustly validated in a chicken model of avian influenza ^9^, demonstrating that *E. coli-*mediated nucleic acid delivery can indeed be used to mitigate the replication and shedding of a clinically important virus.

In this study, we have established our engineered strain, *E. coli* SVC1, as a viable system for in vivo applications, particularly via validation in a murine model. We first characterized the duration of the gene silencing it can mediate in vitro. We then examined the feasibility of directly administering SVC1 to specific tissues and organs and assessed its biodistribution in clinically relevant sites in vivo. Next, we investigated the safety profile of SVC1 upon repeated dosing to the lungs via histological analysis and monitoring of expression level changes in innate and adaptive immunity-related genes. Finally, we validated the ability of SVC1 to deliver therapeutic shRNAs, functioning as an antiviral, using an in vivo influenza virus infection model. Taken together, the results presented here support the potential of SVC1 to serve as a powerful delivery platform for therapeutic nucleic acids and support translational research to drive its future clinical development.

## Results

### *Invasive SVC1* E. coli *cells deliver functional shRNA in vitro*

To assess the RNAi activity of shRNAs expressed from the SiVEC plasmid (pSiVEC) and delivered to eukaryotic cells by invasive, non-pathogenic *E. coli* cells, we measured green fluorescent protein (GFP) depletion in human alveolar basal epithelial cells (A549 cells) stably expressing GFP. The host *E. coli* strain (SVC1) was previously engineered to be invasive to eukaryotic epithelial cells (**Figure 1A**). Via an interaction between *Yersinia pseudotuberculosis* invasin on the bacterial surface and β1 integrin on the surface of the target cells, SVC1 bacteria invade eukaryotic cells via receptor-mediated endocytosis. Upon entry, the bacteria lyse in the endosome to release their cargo, including the shRNA and LLO, the product of *Listeria monocytogenes hlyA*, which perforates the endosome allowing the shRNA to enter the cytoplasm of the invaded cells. We treated A549 cells with SVC1 carrying pSiVEC encoding a GFP shRNA (GFP-shRNA) or pSiVEC encoding a non-targeting small RNA (scramble) for two hours. Subsequently, we removed the bacteria, and the cells were further incubated for an additional 96 hours. We measured GFP expression 6, 24, 48, 72, and 96 hours after bacterial removal (**Figure 1B**). As shown in **Figure 1C**, GFP expression was robustly reduced in A549 cells treated with GFP-shRNA compared with A549 cells treated with scramble. GFP depletion persisted over 96 hours at both a low dose (**Figure 1D**, left) and at a high dose of bacteria (**Figure 1D**, right). As would be expected, the higher dose of invasive bacteria resulted in more robust GFP depletion (i.e., a dose-dependent response), suggesting that the level of depletion achieved via shRNA delivery can be controlled by varying the number of invasive bacteria. The length of the cell cycle of A549 cells under the growth conditions used here is approximately 20 hours (**Supplemental Figure 1**); thus, some cell division likely occurred over the course of this experiment. The observation that GFP expression did not increase during at least the first 72 hours of the time course suggests that the abundance of delivered shRNA is sufficient to be inherited by the daughter cells of the originally invaded cells, which has been previously reported for siRNA ^54^. Taken together, the data presented in **Figure 1** confirm that SVC1 can invade eukaryotic epithelial cells and deliver a cargo of shRNA that is then processed via the RNAi pathway to robustly and persistently silence the expression of a target gene.

**Figure 1.**
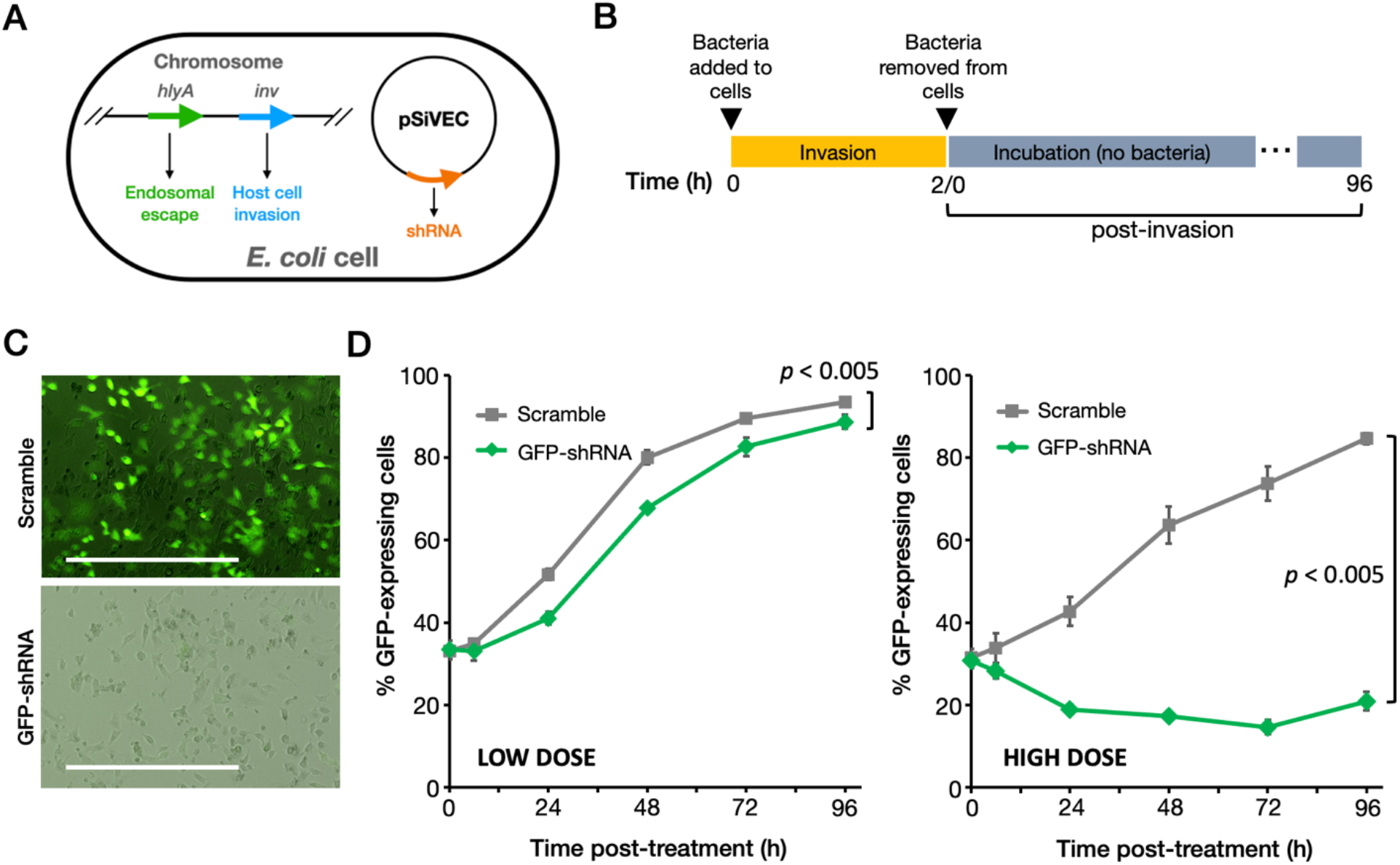
Bacterial delivery of shRNA-GFP silences GFP expression in respiratory epithelial cells. **A –** Genetic features of *E. coli* SVC1 bacteria. **B –** Experimental design for shRNA delivery. A549 cells constitutively expressing a GFP reporter gene were seeded in 24-well culture plates and incubated with two doses of bacteria (low dose, 1.56×10^6^ CFU/mL; high dose, 1×10^8^ CFU/mL) expressing a scramble (non-targeting) small RNA or an shRNA targeting GFP, and GFP expression was monitored for 96 hours post-invasion. **C –** Overlay of GFP and brightfield channels demonstrating GFP knockdown achieved via SVC1-mediated GFP-shRNA delivery (96 hours post-invasion; scale bar represents 1 mm). **D –** GFP knockdown reported as the percentage of GFP-positive A549 cells. Plots show the mean ± SD at each time point. The statistical significance of the differences between the control and experimental groups was assessed using two-way ANOVA, and the p-values are provided.

### *Invasive SVC1* E. coli *can be administered to various epithelial tissue types*

SVC1 binds to target cells via an interaction between its surface-expressed invasin (**Figure 1A**) and β1 integrin on the surface of the target eukaryotic cells. β1 integrin is expressed by epithelial cells in multiple tissue types, including those in the cornea, respiratory tract, reproductive tract, digestive tract, and skeletal muscle. This ubiquity suggests that SVC1 can be used as a flexible approach for delivering therapeutic moieties to various tissues. To examine the versatility of SVC1 as a delivery system, we used various methods (**Table 1**) to apply invasive, fluorescently labelled SVC1 bacteria to the eye, upper (nasal cavity) and lower (lungs) respiratory tract, vagina, digestive tract, and skeletal muscle.

**Table 1.** Routes of administration of invasive SVC1 bacteria to various tissues

| Organ/tissue |  | Administration method | CFU (dose) per volume |
| --- | --- | --- | --- |
| Eye | | Eyedrop | $1 \times 10^9$ CFU / 5 μL |
| Vagina | | Vaginal instillation | $1 \times 10^9$ CFU / 15 μL |
| Skeletal muscle | | Intramuscular (IM) injection, thigh | $1 \times 10^8$ CFU / 25 μL |
| Respiratory tract |  |  |  |
| | Upper (nasal cavity) | Intranasal (IN) | $1 \times 10^9$ CFU / 50 μL |
| | Lower (lungs) | IN droplet inhalation with anesthesia | $1 \times 10^9$ CFU / 50 μL |
| Digestive tract |  |  |  |
| | Oral cavity | Oral gavage | $1 \times 10^9$ CFU / 100 μL |
| | Stomach | Oral gavage | $1 \times 10^9$ CFU / 100 μL |

As shown in **Figure 2A**, SVC1 bacteria remained localized following administration, suggesting that they can indeed be used for organ- and tissue-specific delivery of therapeutic cargo. These results validate the versatility of SVC1 for use as a delivery vehicle in various tissue types. We aim to administer the fewest SVC1 bacterial cells required to achieve the desired effect. While our ongoing work suggests that target cell invasion is efficient, our current routes of in vivo administration likely introduce a surplus of bacteria. In light of this consideration, we were interested in the fate of any excess bacterial cells. The clearance of the system is especially important with regard to trafficking of the SVC1 delivery vehicle to the liver, which is often undesirable in drug delivery applications due to associated toxicity ^55–57^ To explore this feature of SVC1, we administered SVC1 intramuscularly in the hind limb and then collected a section of muscle tissue that received the localized injection, liver, and proximal draining lymph node after 20 and 72 hours. We then tested for the presence of the delivery vehicle in the injected muscle as well as the liver and lymph nodes using PCR with primers that detect the SVC1-borne plasmid (pSiVEC) in total DNA isolated from each tissue sample as shown in the table in **Figure 2B**, the PCR results suggest that the excess bacteria are cleared via the lymphatic system rather than being trafficked to the liver.

**Figure 2.**
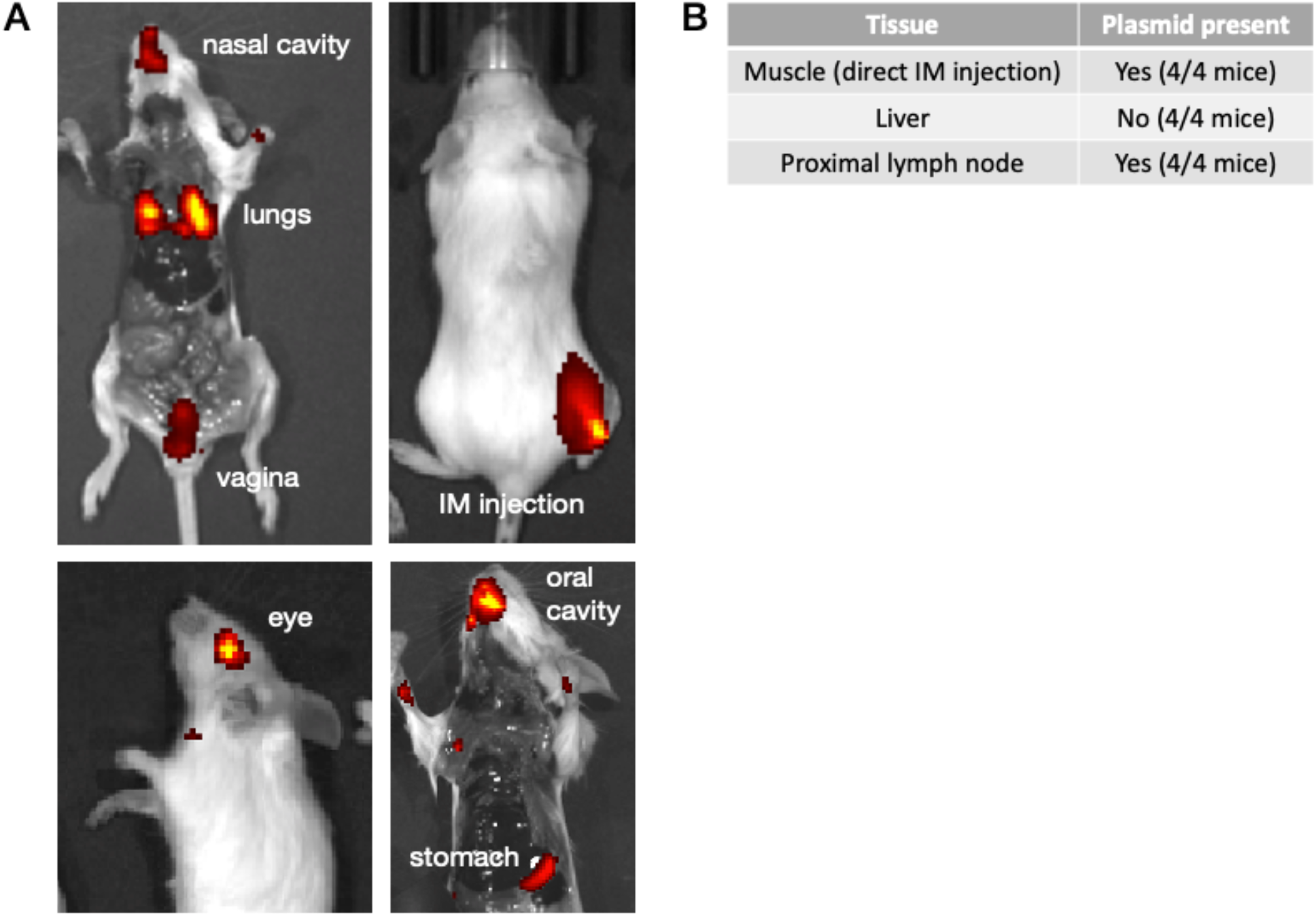
In vivo administration and clearance of invasive bacteria. **A –** Imaging of fluorescently labeled SVC1 bacteria after delivery as described in the text and Table 1. Relative signal intensity is shown ranging from red (low) to yellow (high). **B –** Assessment of plasmid presence in various tissues after IM delivery.

### Repeat dosing of SVC1 to the respiratory tract is well tolerated in vivo

To explore the potential of SVC1 as a tissue-targeting nucleic acid delivery platform, we focused on the respiratory tract. By virtue of their ability to silence expression of target proteins, therapeutic shRNAs have broad usefulness in the treatment of a variety of infectious diseases of the lungs, including respiratory viruses (e.g., influenza and SARS viruses); however, because of the intrinsic instability of shRNAs in the cytoplasm and the transience of the SVC1 bacteria at the delivery site (see above), a robust therapeutic effect would likely require repeat administration. Various issues with repeat dosing (e.g., undesirable immune responses and acquired resistance) limit the usefulness of some current delivery systems, especially adeno-associated virus (AAV)-based systems ^33–37^; therefore, we were interested in whether SVC1 could overcome such limitations, especially detrimental immunogenicity. To this end, we examined the effects of repeat dosing of SVC1 to the respiratory tract. We administered six doses of 50 αL containing 1×10^9^ SVC1 bacteria suspended in sterile phosphate-buffered saline (PBS) intranasally to mice over the course of 60 hours (**Figure 3A**). As a control, we also treated mice with PBS alone (sham). Twenty-four hours after the sixth dose, we euthanized the mice and collected tissue samples for analysis. We monitored posture, grooming, interest in food, and behavior, and all were normal prior to euthanasia, and all of the SVC1-treated mice appeared healthy. Upon dissection, we found no gross abnormalities or differences in the lungs, liver, kidneys, heart, abdominal cavity, nasal cavity, spleen, or other internal anatomy, and we did not observe any gross lesions (data not shown). Nasal sections from bacteria- and sham-treated mice had normal morphology with no inflammation, no bacteria, no increase in mucus secretion, and no alteration in the mucociliary apparatus (representative nasal sections are shown in **Figure 3, B, C**). We also examined the lungs for three pathologies: inflammation, immune cell infiltration, and injury (**Figure 3, D–F**). The lung tissue samples all had minimal to mild increase in alveolar and intracapillary macrophages affecting predominantly one or two lobes that did not vary significantly among the treated and control, suggesting a baseline responsive mild lung infiltrate that was not distinctly due to the treatment nor made worse by the treatment. In rare cases, the blood vessels were minimally reactive and surrounded by edema and rare fibrin, supporting the possibility of a hematogenous antigen source unrelated to either treatment (SVC1 or sham). A single SVC1-treated mouse had severe lymphoplasmacytic and histiocytic pneumonia that varied significantly from the other mice. There was subjectively a moderate lymphoid aggregate (BALT) hyperplasia, supporting a mild baseline immune response; however, this pathology was uniformly present and could not be ascribed to SVC1 bacterial treatment. The immunogenicity of SVC1 was then examined objectively as described in detail below. We did not observe SVC1 bacteria on hematoxylin and eosin-stained sections in any tissue sample, suggesting that the bacteria were rapidly cleared and/or that invasion was rapid and robust. Taken together, these results establish SVC1 as a safe bacterial delivery vehicle for repeated intranasal delivery to the respiratory tissues.

**Figure 3.**
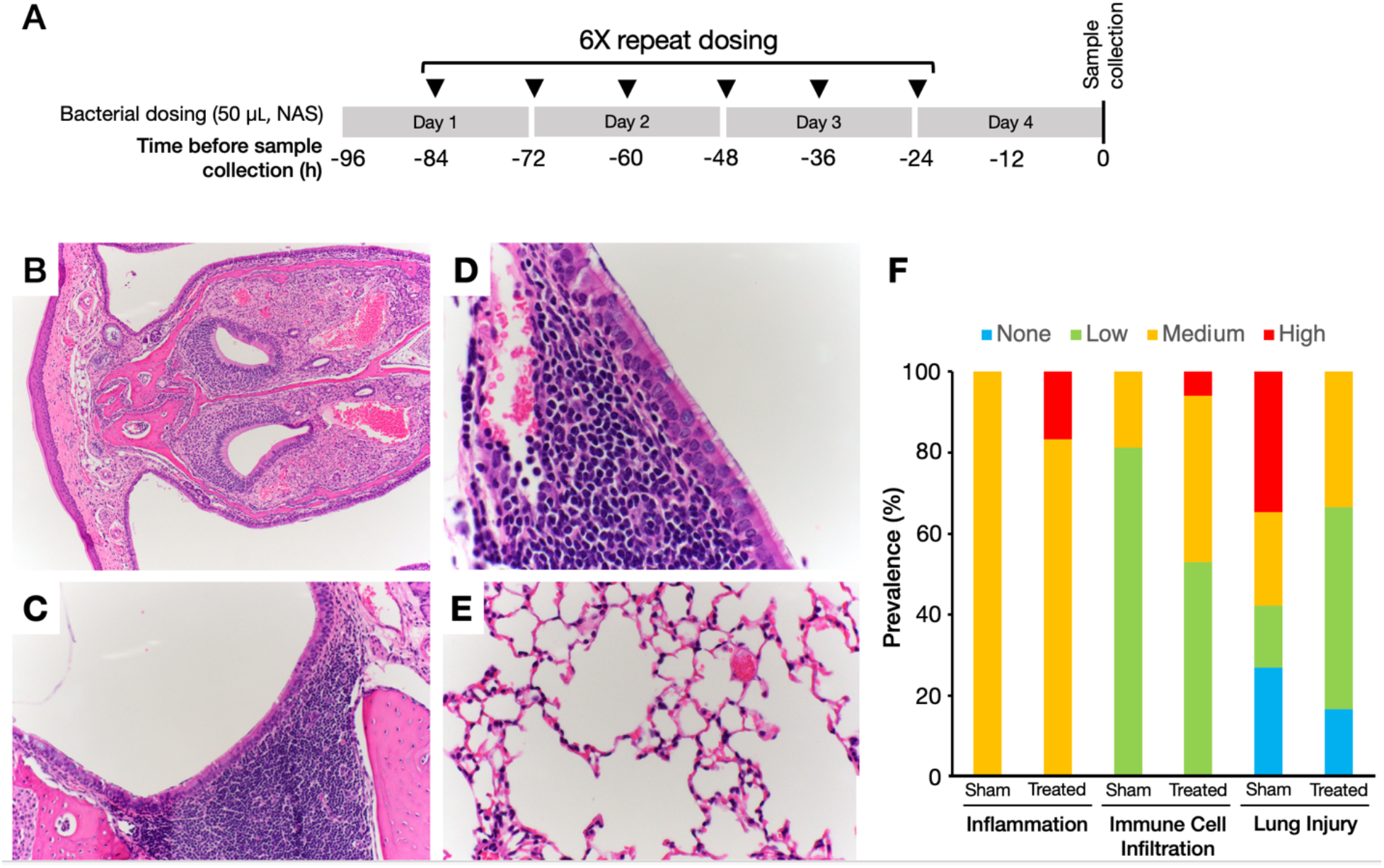
Repeat in vivo dosing of invasive bacteria to the lungs does not adversely affect tissue homeostasis. **A –** Mice were anesthetized and dosed intranasally with invasive bacteria (1×10^8^ CFU per 50 µL dose, n=5) in PBS (treated) or PBS alone (n=5) (sham). Tissue samples were collected for analysis 24 hours after administration of the last dose. **B –** Cross section of the inferior nasal meatus with normal pseudostratified ciliated respiratory epithelium and the vomeronasal organ and glands, **C –** Close-up of normal ciliated respiratory epithelium and underlying nasal mucosa-associated lymphoid tissue, **D –** nasopharyngeal duct and nasal-associated lymphoid tissue, **E –** normal lung tissue. **F –** A summary of the prevalence of inflammation, immune cell infiltration, and injury in the lungs based on histological examination. The categories of low, medium, and high are defined in the *Materials and Methods*.

### SVC1 is minimally immunogenic in the respiratory tract

To evaluate the systemic immunogenicity of SVC1 in the respiratory tract, we collected the spleens from the mice in the repeat dosing experiment, purified total RNA, and examined gene expression differences between sham and SVC1-treated mice using a Qiagen RT^2^ Profiler PCR Array for mouse adaptive and innate immune responses (**Figure 4A**). This array allows simultaneous monitoring of 84 immune-related genes, which taken together are representative of key innate and adaptive immune responses. As SVC1 was developed from a commensal, highly attenuated *E. coli* strain, we expected that it would be minimally immunogenic in the respiratory tract, even after repeated dosing. Among the 84 immune-related genes analyzed, only 7 had statistically significant (α=0.05) expression level changes in the SVC1 bacteria-treated mice compared with the sham-treated mice: *Ccr4, Ccr5, IL1a* (Interleukin 1a), *H2-Q10, Il5* (interleukin 5), *Nlrp3*, and *Rorc* (**Figure 4B**). The upregulation of these genes likely reflects the presence of bacteria, as each has been linked to responses to bacterial lipopolysaccharide (LPS) ^58–65^. The upregulation of LPS-related genes is not unexpected because while the LPS of SVC1 is truncated (lacking the O-antigen), it remains recognizable, though only mildly immunogenic, in mammals ^66–68^. In addition, the gene encoding myeloperoxidase (MPO) was upregulated over 2-fold. While this change was not statistically significant, it could also reflect an effect of bacterial presence as MPO has been linked to a response to pro-inflammatory agents in the lung epithelium ^69^. Repeated treatment did not result in significant non-specific or specific changes in the expression levels of pattern recognition receptors (PRRs), cytokines/chemokines, innate/adaptive markers, inflammatory or bacterial defense markers. Taken together, these data indicate that repeat dosing with SVC1 to the respiratory tract in mice does not induce a robust immune response compared to PBS (sham) dosed mice, suggesting that SVC1 is minimally immunogenic and safe for repeat dosing in a mammal.

**Figure 4.**
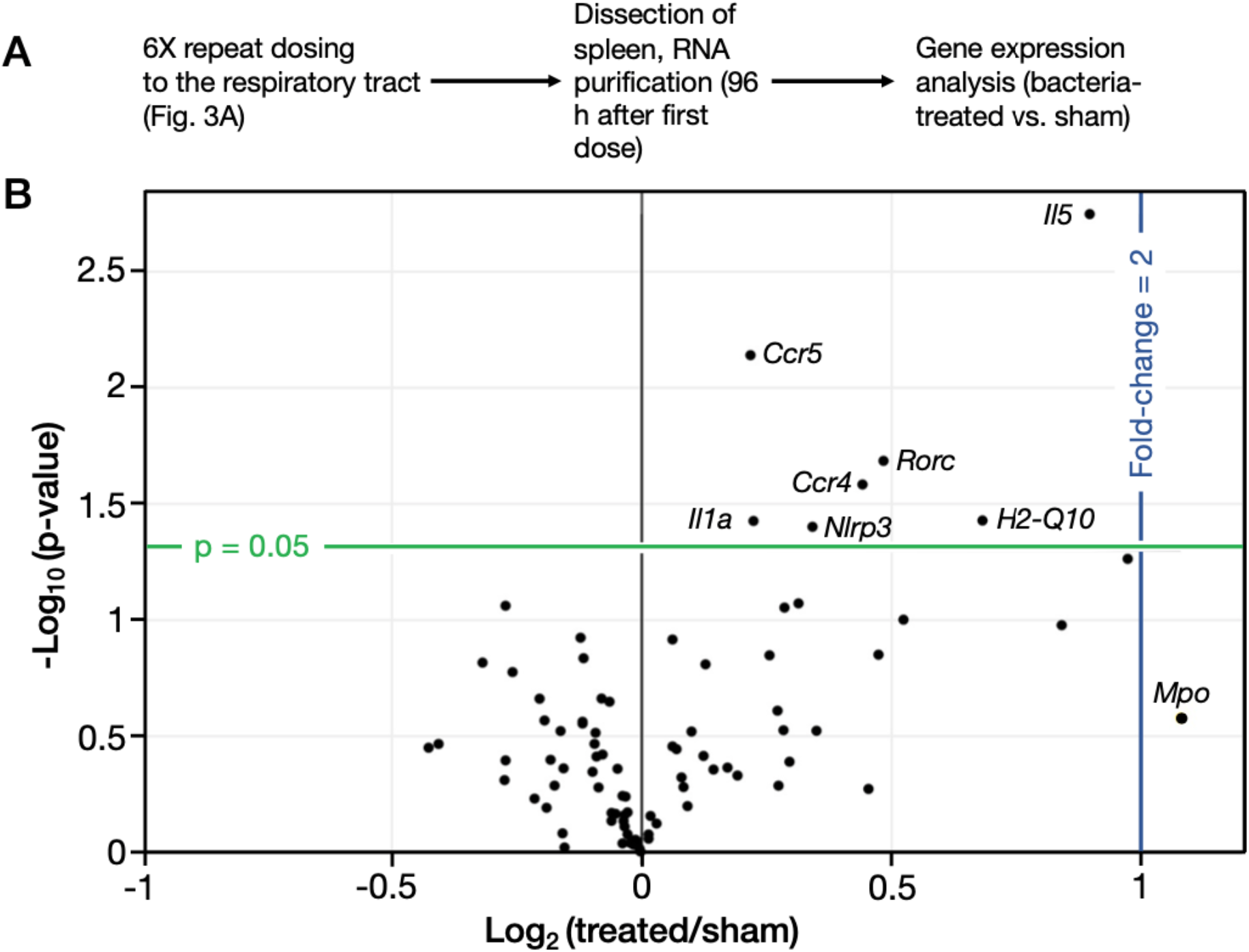
Repeat in vivo dosing of invasive bacteria to the lungs is minimally immunogenic. **A –** Mice were dosed with invasive bacteria or sham as in Fig. 3. The spleens were removed 24 hours after administration of the last dose, and RNA was isolated for analysis using the Qiagen RT^2^ Profiler Mouse Innate and Adaptive Immune Response PCR Array. **B –** A volcano plot of differential gene expression after 6x dosing with invasive bacteria versus sham treatment (PBS). Expression differences were considered statistically significant at p > 0.05 (green line). A fold-change of 2 is indicated by the blue line. The top ten differentially expressed genes and those genes with a statistically significant change in expression are labelled (n=13).

### SVC1 can deliver anti-viral shRNAs to the respiratory tract in vivo

Finally, we wanted to demonstrate the in vivo therapeutic potential of SVC1 to deliver anti-viral shRNAs. To this end, we designed two therapeutic strains: an SVC1 derivative expressing an shRNA against the influenza A virus (IAV) PA protein (RNA polymerase complex subunit) (SVC1-PA) and an SVC1 derivative expressing an shRNA against the influenza NP protein (nucleocapsid) (SVC1-NP). These strains are mixed prior to administration to produce the SiVEC-IAV cocktail. Upon simultaneous delivery of the shRNAs to the cytoplasm of a respiratory epithelial cell (the site of IAV replication) via administration of SiVEC-IAV, they are processed via the RNAi pathway into siRNAs that silence PA and NP expression, thereby inhibiting IAV replication and reducing viral shedding. To test the efficacy of these shRNAs delivered via SVC1, we dosed mice with a cocktail of SVC1-PA and SVC1-NP at three doses (low, medium, high) twice prior to viral challenge and then four times after the mice were exposed to H1N1 IAV (PR8 strain) as described in **Figure 5A**. On days 3, 5, 7, and 9 post-challenge, we collected the nasal turbinates, purified total RNA, and determined the viral titers (as EID50 equivalent/mL) via reverse transcription quantitative PCR (RT-qPCR). As shown in **Figure 5B**, reductions in viral titer were observed at the low, medium, and high bacterial doses, with a clear dose-response trend. As expected, the highest dose was the most effective at reducing viral titer in the nasal turbinates. These results demonstrate that SVC1 can be used as an effective vehicle for the delivery of therapeutic shRNAs to the lungs in a respiratory disease model.

**Figure 5.**
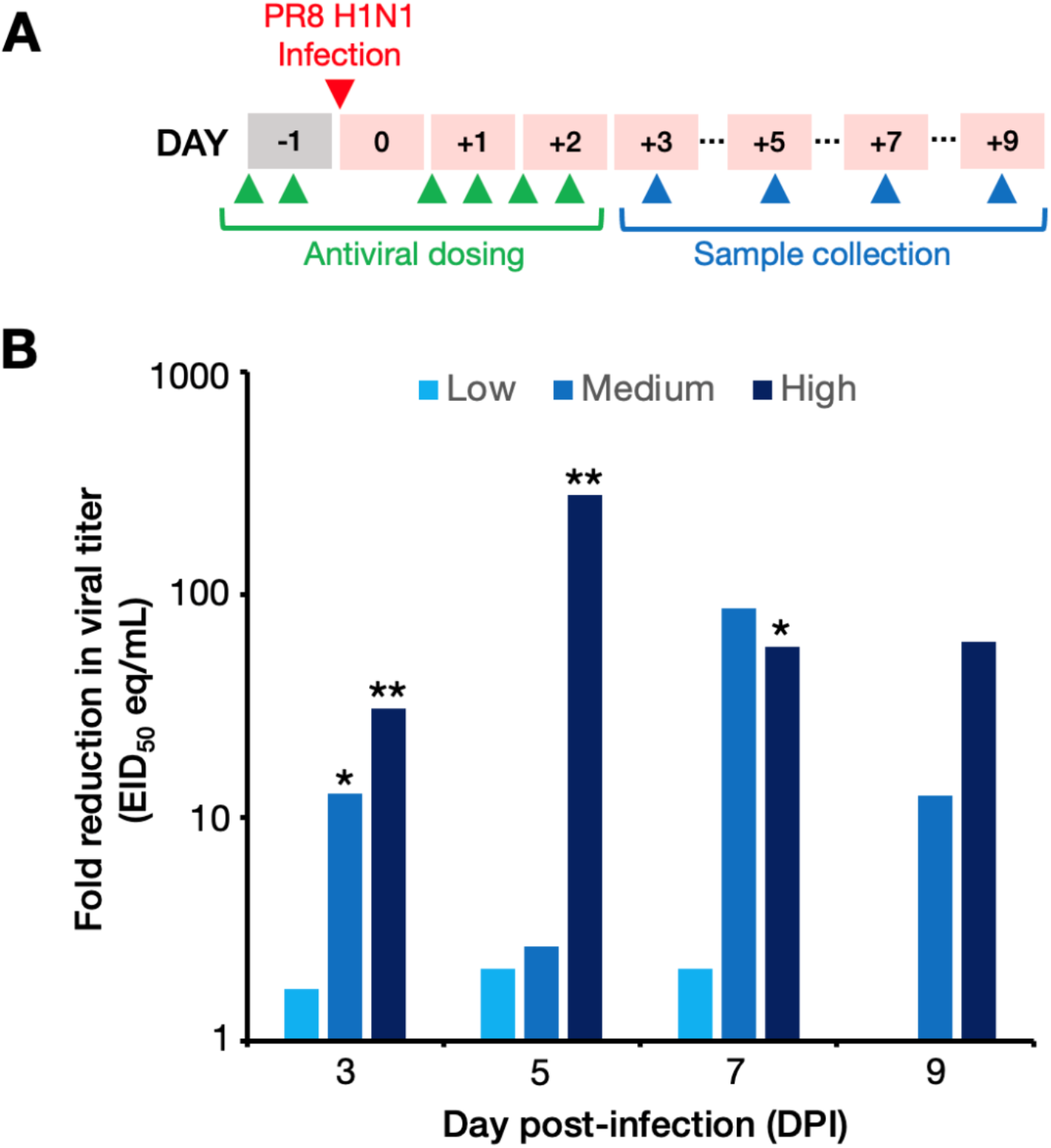
Bacterially delivered influenza-targeting shRNAs mitigate influenza virus replication in vivo. **A –** Mice (n=6 per group) were treated with high (1×10^8^ CFU/mL), medium (1×10^7^ CFU/mL), or low (1×10^6^ CFU/mL) doses of bacteria expressing influenza-targeting shRNAs or PBS sham twice prior to and four times after (green arrows) infection with PR8 (H1N1) influenza virus (red arrow). Samples of nasal turbinate tissue were collected for assessment of viral titer up to nine days post-infection. **B –** Plot showing the fold reduction in viral titer (as EID50 eq/mL, see *Materials and Methods*) in mice dosed with shRNA-expressing bacteria or PBS (sham) from three to nine days post-infection (DPI). Statistical significance was calculated using the two-sample Wilcoxon rank-sum (Mann-Whitney) test. Asterisks indicate statistical significance: *, p ≤ 0.05; **, p ≤ 0.01.

## Discussion

A significant factor limiting the translation of therapeutic nucleic acids, proteins, and gene editing technologies from bench to bedside is the absence of safe and robust vehicles for targeted delivery to affected cells and tissues. In the case of nucleic acids, this limitation is imposed by the negative charge, instability, immunogenicity, and in some cases the large size of nucleic acids, particularly relative to typical small-molecule drugs. The SVC1-based platform described here offers an elegant solution to the targeted delivery conundrum and holds promise for utility in ameliorating a range of diseases and disorders. SVC1 was engineered 1) to constitutively express nucleic acids, 2) to target clinically relevant cell types (i.e., mucosal epithelial cells), and 3) to escape the endosome allowing the release of the nucleic acids into the target cell cytoplasm. SVC1 can be tailored to produce different types of therapeutically relevant nucleic acids. Furthermore, based on the targeted administration results shown in **Figure 2**, SVC1 might be useful for treating a plethora of oral, respiratory, gastrointestinal, ocular, vaginal, and rectal diseases. While not investigated here, SVC1 can be used for delivery of proteins, eukaryote-translatable mRNA, and gene-editing systems (e.g., CRISPR/Cas) to targeted mucosal tissues, providing cellular uptake and, where appropriate, nuclear translocation of the gene-editing nuclease system, without the need for host genome integration.

A key feature of any drug delivery system is the amount of therapeutic moiety that can be delivered to the targeted cells. In human cells, siRNAs are not enzymatically replicated to allow transgenerational inheritance of the silencing effect, in contrast to the well-known transgenerational inheritance of RNAi in nematodes ^70^; therefore, delivered shRNAs are diluted during cell division as the finite supply of molecules is depleted upon partitioning to daughter cells. While it was not technically feasible to measure the number of shRNAs delivered by SVC1 bacteria at the time this work was completed, the persistence of the silencing effect shown in **Figure 1D** provides hints to understanding this key feature of the system. The unexpected finding that the silencing effect of the shRNAs produced and delivered by SVC1 were sustained for at least 96 hours post-treatment, a time period sufficient for the initially invaded cells to divide ^71,72^, suggests that SVC1 cells deliver a large payload of shRNAs, attesting to the potential for a sustained therapeutic effect. Importantly, this effect is achieved without genome integration and is ultimately transient as the populations of shRNAs and siRNAs are depleted either via dilution upon cell division or due to the inherent instability of RNA molecules (the latter being especially relevant to the transiency of the effect in post-mitotic cells).

The intracellular delivery modalities currently in use in the clinic, e.g., lipid nanoparticles and viral vectors, suffer from undesirable immune effects that can limit their utility, particularly for repeated dosing. The preliminary analysis of the immunogenicity of SVC1 delivered to the lungs (after 6 repeated doses over three days) presented here suggests that the bacteria are minimally immunogenic, as no significant immune cell infiltration (**Figure 3F**) nor statistically significant changes in the expression level of any screened immune-related gene (2-fold or greater) (**Figure 4**) were observed. Interestingly, of the systemically upregulated genes (n=7) detected after repeated respiratory administration, genes related to cellular responses to LPS were overrepresented (7 out of 7) ^58–65^. This observation suggests that the array analysis used here is sufficiently sensitive to detect subtle gene expression changes in response to the presence of the bacterial LPS but that the amount of LPS delivered via this administration scheme was sufficiently low to not induce a robust systemic immune response. Finally, tissue damage was not observed even in tissue directly exposed to the bacterial vehicle (**Figure 3B–E**). We are currently further modifying SVC1 to express a less-immunogenic LPS to further mitigate immunogenicity concerns. Work is also underway to explore whether the SVC1 system is affected by acquired immunity. However, an acquired immune response to the SVC1 bacteria is not anticipated based on the gene expression analysis described here as well as the additional genetic modifications we are making to further attenuate the LPS of SVC1.

To demonstrate the potential of SVC1 as an in vivo therapeutic delivery vehicle, we demonstrated that simultaneous delivery of shRNAs designed to silence two essential influenza genes (the SiVEC-IAV cocktail) could ameliorate viral replication in a mouse model of influenza infection (**Figure 5**). To our knowledge, this work represents the first demonstration of the use of a bacteria-based delivery system in a mammalian antiviral application. The robust reductions in viral replication shown in **Figure 5B** confirm that SVC1 can indeed be used to deliver therapeutic RNA molecules. Importantly, our data also revealed a dose-dependent reduction in viral replication when different numbers of SVC1 cells were intranasally administered. This dose responsiveness demonstrates that the number of bacteria delivered can be modulated to achieve different therapeutic outcomes, which might be advantageous in some applications of such a platform (e.g., delivery of gene editing components).

Finally, a distinguishing advantage of the SVC1 bacterial delivery platform (in comparison to other available delivery platforms) is that the bacteria themselves can produce the therapeutic moieties that they deliver, as demonstrated here by the bacterial transcription of shRNAs that feed into the host cellular RNAi pathway. This feature of the system eliminates RNA manufacturing steps and production costs ^73–76^. As it is simple and fast to generate large quantities of bacteria using widely available manufacturing approaches, SVC1-based therapeutic products could be readily generated in massive quantities from a small stock, and if properly stored, could have a long shelf life. With manufacturing in mind, we are currently working on characterization of potency, including developing methods to quantitate the number of shRNA molecules generated per SVC1 bacterial cell.

Our ongoing research and development efforts are focused on optimization of SVC1 as a platform for the production (via bacterial transcription) and delivery of both linear and circular eukaryote-translatable mRNAs and for the production and delivery of gene editing proteins and RNAs (i.e., CRISPR/Cas machinery). Due to the vast genetic coding capacity and transcriptional flexibility of *E. coli*, SVC1 can express and deliver high molecular weight RNA molecules. Furthermore, our vast knowledge of *E. coli* molecular genetics enables further application-specific optimization (e.g., additional modulation of RNase activities) to improve its performance as a highly versatile delivery platform. We expect that the advantages offered by live bacteria, and SVC1 in particular, will lead to future studies that further enable and validate the usefulness of bacteria as a powerful multi-application delivery platform.

## Materials and Methods

### Bacterial strains, plasmids, and cells

The invasive *E. coli* strain SVC1 is a K-12 derivative [F^−^ *endA1 hsdR17 (rK*^*–*^ *mK*^*+*^*) glnV44 thi-1 relA1 rfbD1 spoT1* Δ*rnc* Δ*dapA*]. The cells are auxotrophic for diaminopimelic acid (DAP) due to a deletion of *dapA. E. coli* cells were cultured in brain-heart infusion (BHI) medium supplemented with DAP (100 µg/mL) and appropriate antibiotics at the following concentrations: kanamycin, 25 µg/mL; ampicillin, 100 µg/mL. A549 cells are a human adenocarcinoma alveolar basal epithelial cell line. For the in vitro GFP silencing experiments, an A549 cell line constitutively expressing GFP (Cell Biolabs, San Diego, California, AKR-209) was used. The identity of the A549/GFP cells was confirmed by GFP expression, morphology, and trypan-blue dye exclusion, and all cell cultures were routinely monitored for microbial contamination using standard techniques. The construction of pSiVEC-scramble (non-specific small RNA sequence), pSiVEC-PA, and pSiVEC-NP, which were derived from pmbv43 ^8^, is described in detail elsewhere ^9^. pSiVEC-GFP was constructed using the DNA template encoding the shRNA specific for GFP ^77^ from the copepod *Pontellina plumata*: sense, GCTACGGCTTCTACCACTTT and antisense, AAAGTGGTAGAAGCCGTAGC. Using standard cloning and transformation methods, resulting SVC1 colonies transformed with the plasmids pSiVEC-scramble, pSiVEC-PA, pSiVEC-NP, and pSiVEC-GFP were screened by PCR, and a single positive clone was sequence validated and propagated. Stocks were generated and stored at -80 °C in 20% glycerol. A single frozen aliquot from each construct stock was thawed to determine colony forming units (CFU)/mL via plate enumeration ^9^. These strains are referred to as SVC1-scramble, SVC1-PA, SVC1-NP, and SVC1-GFP.

#### In vitro invasion assay

For the GFP gene silencing studies (**Figure 1**), A549/GFP cells were seeded one day prior to invasion in a 24-well tissue culture-treated microplate with black walls and clear bottom (Perkin Elmer, Waltham, MA; VisiPlate 1450) to allow the monolayer to reach 70% confluence. On the day of invasion, 1-mL frozen aliquots of SVC1-GFP and SVC1-scramble were thawed, centrifuged, re-suspended and appropriately diluted in DMEM/DAP. A549/GFP cells were washed to remove antibiotics and incubated with SVC1-GFP or SVC1 -scramble at a low dose (1.56×10^6^ CFU/mL) and high dose (1×10^8^ CFU /mL) at 37 °C with 5% CO2. After 2 hours incubation, A549/GFP cells were washed three times, and fresh DMEM with antibiotics was added. GFP signal was measured at 0, 6, 24, 48, 72, and 96 hours post invasion using the Nexcelom Celigo (Lawrence, MA) and reported as the percentage of GFP-positive A549 cells. The statistical significance of the differences between the SVC1-scramble control and SVC1 - GFP experimental groups was assessed using two-way ANOVA (p < 0.005).

#### In vivo biodistribution assays

To characterize the biodistribution of the SiVEC vehicle following localized administration to mouse mucosal epithelial and skeletal muscle tissues (**Figure 2A**) SVC1-scramble and isogenic non-invasive SVC1 bacteria (lacking pSiVEC-scramble) were fluorescently labeled using the XenoLight RediJect 750 near-infrared fluorescent probe (Perkin Elmer, Waltham, MA). Nine-week-old female BALB/c mice (Jackson Laboratory, Bar Harbor, ME) were anesthetized with inhaled isoflurane in an anesthesia chamber and then transferred to the Perkin Elmer IVIS Spectrum system for in vivo imaging. Mice were treated with SVC1-scramble (invasive) and an untreated control mouse was included for calibration of background signal due to autofluorescence. The route of administration and dose per tissue type is shown in Table 1.

To demonstrate localized delivery to mucosal epithelia (**Figure 2A**), mice were imaged at 0, 2, 4, 6, and 18 hours post-administration to the eyes, lungs, nose, vagina, oral cavity, and stomach as described in Table 1. Eighteen hours post-treatment, the mice were euthanized, and the body cavity of each animal was opened with a ventral, longitudinal incision, extending from the vaginal opening, up through the lower jaw to expose deeper tissues difficult to image. One final image was captured of all mice.

To demonstrate injected delivery to the skeletal muscle (thigh) and to characterize the route of bacterial vehicle clearance (**Figure 2B**), mice were imaged at 0, 6, 20, 48, and 72 hours post-IM injection. 72 hours post-IM injection, mice (n=4) treated with SVC1-scramble were euthanized, and liver, draining lymph tissue, and the thigh muscle were collected for subsequent DNA extraction to screen for the presence of SVC1-scramble. Briefly, DNA was extracted from 20 mg of each tissue using the ZYMO Quick-DNA™ Miniprep Plus Kit (Zymo Research, Irvine, CA). Resulting DNA was amplified neat and diluted 1:2, and 1:5 in molecular water with conventional PCR using primers specific to pSiVEC-scramble: forward, CAGATGCGTAAGGAGAAAATACCGCAT; reverse, CATTAATGAATCGGCCAACGCGCG. PCR amplification was completed using a 25 µL reaction containing 5 µL DNA template, 12.5 µL 10X GoTaq Master Mix (Promega Corporation, Madison, WI), and 1 µM final each primer. Cycling conditions consisted of 94 °C for 4 minutes followed by 30 cycles of 94 °C for 30 seconds, 56 °C for 30 seconds, and 72 °C for 2 minutes, and a final elongation step at 72 °C for 10 minutes. PCR products were analyzed by 2% agarose gel electrophoresis. The limit of detection for DNA extraction and PCR amplification of SVC1-scramble was 10^2^ CFU/20 mg tissue.

#### Immunogenicity assay and histopathology

The safety of repeat dose administration to the lungs and the effect on tissue pathology and immune response was evaluated in 9-week-old female BALB/c mice. SVC1-scramble or a PBS-sham treatment was administered intranasally to n=5 mice per group approximately every 12 hours for six doses total (**Figure 3A**). 24 hours after administration of the last dose, mice were euthanized for necropsy and gross pathology assessment and the collection of tissues for histopathology and gene expression profiling. The spleen was collected from each animal and stored in RNAlater (Thermo Fisher Scientific, Waltham, MA) for subsequent RNA extraction and analysis using the Qiagen RT^2^ Profiler Mouse Innate and Adaptive Immune Response PCR Array (Qiagen, Hilden, Germany). The lungs were placed in a tissue cassette, and fixed in 10 volumes of 10% neutral buffered formalin (NBF) for 24 hours prior to submission to the histology division of the Colorado State University Veterinary Diagnostic Laboratory (CSU VDL). The head was severed, and the lower jaw, skin and, excess tissue, including cheek fat pads and muscle, were removed. The posterior/rostral two thirds of the skull and brain were removed, exposing the olfactory bulb, but leaving the skull on the lateral aspects of the head. This portion of the head containing the nasal cavity was immediately fixed in 10-volumes of 10% NBF for 24 hours at room temperature. After 24 hours, the fixed heads were rinsed 5x with deionized water, decalcified in 25 mL of 5% formic acid with moderate agitation for 24 hours at room temperature, and rinsed 3x with 0.01 M PBS. The heads were then cut into three sections to expose the nasal cavity and nasal turbinate structure, arranged in a tissue cassette, and transferred back into 10% NBF to fully cover each cassette. The head samples were then submitted to the CSU VDL. Fixed lung and head samples were embedded in paraffin, serially sectioned, mounted, and stained with hematoxylin and eosin (H&E). Slides were pathologically evaluated using a five-point scale to score the degree of inflammation (fibrin, edema, vasculitis, bacteria presence), immune cell infiltration (neutrophils, lymphocytes, plasma cells, macrophages), and lung injury (bronchial epithelial hyperplasia, emphysema). The following scale was used to score each slide: 0: absent (“none”), 1: minimal (“low”), 2–3: mild to moderate (“medium”), and 4: severe (“high”).

Total RNA was extracted from 5 mg of spleen lysate using the Omega Biotek Total RNA 96 kit (Omega Bio-Tek, Norcross, GA) and a KingFisher Flex Instrument (ThermoFisher Scientific, Waltham, MA), and RNA concentration and purity was determined via Nanodrop (ThermoFisher Scientific) spectrometry. The Qiagen RT^2^ First Strand Kit was used to reverse-transcribe 500 ng of each spleen RNA sample, per kit instructions. The resulting cDNA was stored at -20 °C until analysis on the immune array. Expression of 84 immune-related genes was analyzed by qPCR with the Qiagen Mouse Innate & Adaptive Immune Response RT^2^ Profiler PCR Array, per kit instructions. A single sample (one mouse) was assessed on each array plate on a Roche Light Cycler 480 II real time PCR thermal cycler (Roche, Basel, Switzerland). All data were analyzed with Qiagen’s Gene Globe online analysis application. Gene expression was normalized to four housekeeping genes: beta-actin, beta-glucuronidase, heat shock protein 90 (alpha), and beta-2 microglobulin. Gene expression changes were considered statistically significant at p > 0.05, while functionally significant differences were defined as a fold increase of ≥ 2. Results were independently reviewed by a qualified immunologist (subject matter expert) for assessment of a biologically relevant immune response to the SVC1 bacterial treatment.

#### In vivo IAV challenge assays

To demonstrate the therapeutic potential of the SiVEC delivery vehicle, SVC1-PA and SVC1-NP were constructed to express shRNAs targeting the influenza A viral PA and NP mRNAs, respectively, for delivery to the respiratory tissues in an established murine influenza disease model ^78–80^. These strains were generated as previously described ^9^ and were mixed 1:2 (SVC1-PA:SVC1-NP) to create an antiviral cocktail referred to as SiVEC-IAV. Eight-week-old female BALB/c mice (n=200) were anesthetized with inhaled isoflurane and dosed with SiVEC-IAV or PBS-sham by intranasal instillation. Mice were treated with 50 µL of PBS or SiVEC-IAV in high (1×10^8^ CFU/mL), medium (1×10^7^ CFU/mL), or low (1×10^6^ CFU/mL) doses twice prior to and four times after infection with 1×10^6^ EID50 per 50 µL dose of influenza A virus, A/Puerto Rico/89VMC3/1934 (H1N1) (BEI Resources, NIAID, NIH, NR-29028) (see **Figure 5A**). Three, 5, 7, and 9 days post infection (DPI), mice (n=6 per treatment/dose and time point) were euthanized, and nasal turbinates were collected and placed in RNALater.

H1N1 virus titers in the nasal turbinates of mice treated with the high, medium, and low SiVEC-IAV doses were measured via RT-qPCR and fold-reductions in viral titer were calculated relative to the PBS-sham control group. Briefly, total RNA was extracted from approximately 5 mg of nasal turbinate tissue using the Omega Mag-Bind® Total RNA 96 RNA extraction kit with a KingFisher Flex purification system, and RNA concentration and purity was determined using a Nanodrop system. RNA was diluted to 3 ng/µL and reverse transcribed and amplified using the Power SYBR Green RNA-to-CT 1-Step Kit (Thermo Fisher Scientific) to detect the presence of influenza A matrix gene (M-gene). Primer sequences were as follows: M-gene forward, CTTCTAACCGAGGTCGAAACGTA, and M-gene reverse primer, GGTGACAGGATTGGTCTTGTCTTTA. RT-qPCR amplification was completed using a 20 µL reaction containing 5 µL RNA template, 10 µL 2X Power SYBR Green, 0.16 µL 125X RT enzyme, and 0.2 µM of each primer. The RT-qPCR step conditions were 48 °C for 30 minutes, 95 °C for 10 minutes, 40 cycles of 95 °C for 15 seconds, and 60 °C for 1 minute, followed by 60 °C for 5 seconds and 95 °C for 5 seconds to visualize the melting curve for each RT-qPCR assay. The standard curve for virus quantification was generated in triplicate using a series of 10-fold dilutions from 1×10^1^ to 1×10^10^ of the H1N1 stock virus from which the EID50 equivalent per mL (EID50 eq/mL) of each sample was calculated. The limit of detection was determined to be 10^1^ EID50/ml (1 log10 EID50/ml) per reaction. Statistical significance in fold reduction in viral titer between treated and PBS sham mice was calculated using the two-sample Wilcoxon rank-sum (Mann-Whitney) test (p < 0.05).

## Acknowledgements

We thank Alan Schenkel for helpful discussions of the immune response data and Stephanie Morphet-Tepp for technical assistance. The research reported in this study was supported by *the National Center for Allergy and Infectious Disease* of the National Institutes of Health under award number 1R43AI140243-01A1.

---

**Supplemental Figure 1.**
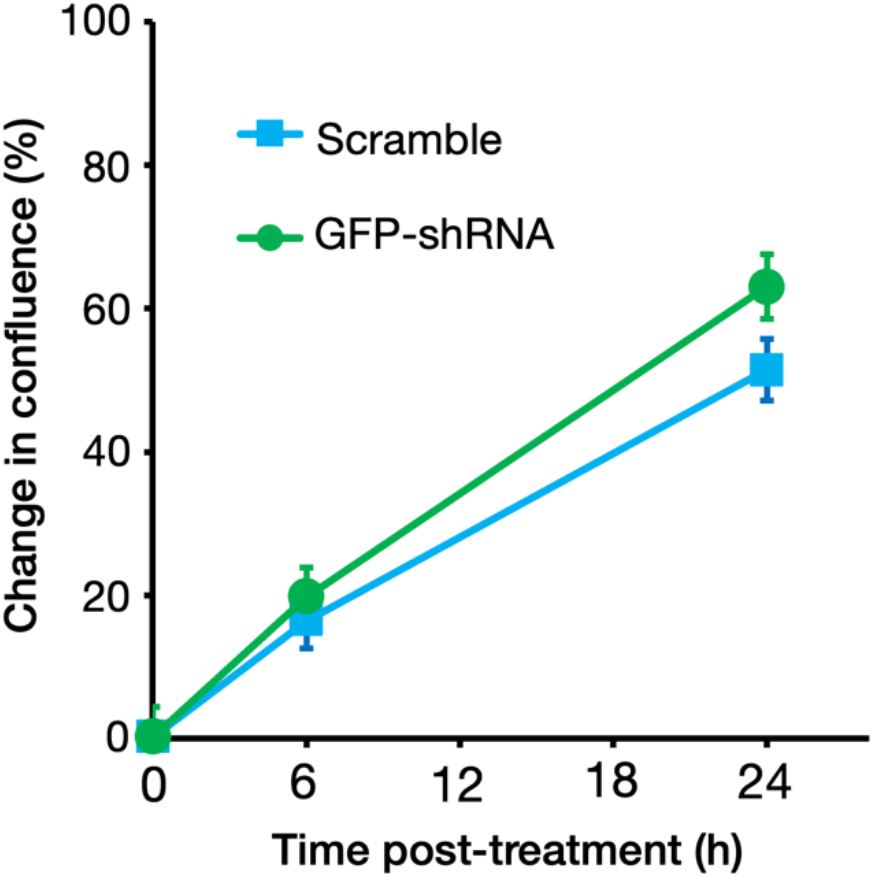
Proliferation of A549 cells under the growth conditions described in the Materials and Methods. Plot shows the mean ± SD at each time point.

## References

1 Hammond SM, Aartsma-Rus A, Alves S, Borgos SE, Buijsen RAM, Collin RWJ, et al. Delivery of oligonucleotide-based therapeutics: challenges and opportunities. EMBO Mol Med 2021. https://doi.org/10.15252/emmm.202013243.

2 Xu L, Anchordoquy T. Drug delivery trends in clinical trials and translational medicine: Challenges and opportunities in the delivery of nucleic acid-based therapeutics. J Pharm Sci 2011. https://doi.org/10.1002/jps.22243.

3 Aguilar ZP. Chapter 5 - Targeted Drug Delivery. Nanomater. Med. Appl. 2013.

4 Manzari MT, Shamay Y, Kiguchi H, Rosen N, Scaltriti M, Heller DA. Targeted drug delivery strategies for precision medicines. Nat Rev Mater 2021. https://doi.org/10.1038/s41578-020-00269-6.

5 Gardlik R, Fruehauf JH. Bacterial vectors and delivery systems in cancer therapy. IDrugs 2010.

6 Fajac I, Grosse S, Collombet JM, Thevenot G, Goussard S, Danel C, et al. Recombinant Escherichia coli as a gene delivery vector into airway epithelial cells. J Control Release 2004. https://doi.org/10.1016/j.jconrel.2004.03.025.

7 Larsen MDB, Griesenbach U, Goussard S, Gruenert DC, Geddes DM, Scheule RK, et al. Bactofection of lung epithelial cells in vitro and in vivo using a genetically modified Escherichia coli. Gene Ther 2008. https://doi.org/10.1038/sj.gt.3303090.

8 Buttaro C, Fruehauf J. Engineered E. coli as Vehicles for Targeted Therapeutics. Curr Gene Ther 2010. https://doi.org/10.2174/156652310790945593.

9 Linke LM, Wilusz J, Pabilonia KL, Fruehauf J, Magnuson R, Olea-Popelka F, et al. Inhibiting avian influenza virus shedding using a novel RNAi antiviral vector technology: proof of concept in an avian cell model. AMB Express 2016. https://doi.org/10.1186/s13568-016-0187-y.

10 Hwang L, Fein SH, Ng K, D’Cruz O, Qazi S, Wang W, et al. RNAi-mediated beta-catenin knockdown in the gastrointestinal mucosa (GI) of patients with familial adenomatous polyposis (FAP): Results of START-FAP trial. J Clin Oncol 2017. https://doi.org/10.1200/jco.2017.35.15_suppl.e15065.

11 Trieu V, Hwang L, Ng K, D’Cruz O, Qazi S, Fong A. First-in-human Phase I study of bacterial RNA interference therapeutic CEQ508 in patients with familial adenomatous polyposis (FAP). Ann Oncol 2017. https://doi.org/10.1093/annonc/mdx393.041.

12 Grillot-Courvalin C, Goussard S, Courvalin P. Bacteria as gene delivery vectors for mammalian cells. Curr Opin Biotechnol 1999. https://doi.org/10.1016/S0958-1669(99)00013-0.

13 Grillot-Courvalin C, Goussard S, Huetz F, Ojcius DM, Courvalin P. Functional gene transfer from intracellular bacteria to mammalian cells. Nat Biotechnol 1998. https://doi.org/10.1038/nbt0998-862.

14 Uribe-Querol E, Rosales C. Phagocytosis: Our Current Understanding of a Universal Biological Process. Front Immunol 2020. https://doi.org/10.3389/fimmu.2020.01066.

15 Isberg RR, Leong JM. Multiple β1 chain integrins are receptors for invasin, a protein that promotes bacterial penetration into mammalian cells. Cell 1990. https://doi.org/10.1016/0092-8674(90)90099-Z.

16 Isberg RR, Barnes P. Subversion of integrins by enteropathogenic Yersinia. J Cell Sci 2001. https://doi.org/10.1242/jcs.114.1.21.

17 Conte MP, Longhi C, Buonfiglio V, Polidoro M, Seganti L, Valenti P. The effect of iron on the invasiveness of Escherichia coli carrying the inv gene of Yersinia pseudotuberculosis. J Med Microbiol 1994. https://doi.org/10.1099/00222615-40-4-236.

18 Humphries JD, Byron A, Humphries MJ. Integrin ligands at a glance. J Cell Sci 2006. https://doi.org/10.1242/jcs.03098.

19 Barczyk M, Carracedo S, Gullberg D. Integrins. Cell Tissue Res 2010. https://doi.org/10.1007/s00441-009-0834-6.

20 Beaulieu JF. Integrins and human intestinal cell functions. Front Biosci 1999. https://doi.org/10.2741/a429.

21 Sheppard D. Functions of pulmonary epithelial integrins: From development to disease. Physiol Rev 2003. https://doi.org/10.1152/physrev.00033.2002.

22 McKay TB, Schlötzer-Schrehardt U, Pal-Ghosh S, Stepp MA. Integrin: Basement membrane adhesion by corneal epithelial and endothelial cells. Exp Eye Res 2020. https://doi.org/10.1016/j.exer.2020.108138.

23 Blandin AF, Renner G, Lehmann M, Lelong-Rebel I, Martin S, Dontenwill M. ß 1 integrins as therapeutic targets to disrupt hallmarks of cancer. Front Pharmacol 2015. https://doi.org/10.3389/fphar.2015.00279.

24 Cooper J, Giancotti FG. Integrin Signaling in Cancer: Mechanotransduction, Stemness, Epithelial Plasticity, and Therapeutic Resistance. Cancer Cell 2019. https://doi.org/10.1016/j.ccell.2019.01.007.

25 Curtiss R. Biological containment and cloning vector transmissibility. J Infect Dis 1978. https://doi.org/10.1093/infdis/137.5.668.

26 Osborne SE, Brumell JH. Listeriolysin O: From bazooka to Swiss army knife. Philos Trans R Soc B Biol Sci 2017. https://doi.org/10.1098/rstb.2016.0222.

27 Mathew E, Hardee GE, Bennett CF, Lee KD. Cytosolic delivery of antisense oligonucleotides by listeriolysin O-containing liposomes. Gene Ther 2003. https://doi.org/10.1038/sj.gt.3301966.

28 Xiang S, Fruehauf J, Li CJ. Short hairpin RNA-expressing bacteria elicit RNA interference in mammals. Nat Biotechnol 2006. https://doi.org/10.1038/nbt1211.

29 Bachmann BJ. Derivations and genotypes of some mutant derivatives of Escherichia coli K-12. Escherichia Coli Salmonella Cell Mol Biol 2nd EdASM Press Washington, DC 1996.

30 Borruat G, Roten CAH, Fay LB, Karamata D. A high-performance liquid chromatography method for the detection of diaminopimelic acid in urine. Anal Biochem 2001. https://doi.org/10.1006/abio.2001.5016.

31 Emamalipour M, Seidi K, Zununi Vahed S, Jahanban-Esfahlan A, Jaymand M, Majdi H, et al. Horizontal Gene Transfer: From Evolutionary Flexibility to Disease Progression. Front Cell Dev Biol 2020. https://doi.org/10.3389/fcell.2020.00229.

32 Meselson M, Yuan R. DNA restriction enzyme from E. coli. Nature 1968. https://doi.org/10.1038/2171110a0.

33 Mingozzi F, High KA. Immune responses to AAV vectors: Overcoming barriers to successful gene therapy. Blood 2013. https://doi.org/10.1182/blood-2013-01-306647.

34 van Haasteren J, Hyde SC, Gill DR. Lessons learned from lung and liver in-vivo gene therapy: implications for the future. Expert Opin Biol Ther 2018. https://doi.org/10.1080/14712598.2018.1506761.

35 Nidetz NF, McGee MC, Tse L V., Li C, Cong L, Li Y, et al. Adeno-associated viral vector-mediated immune responses: Understanding barriers to gene delivery. Pharmacol Ther 2020. https://doi.org/10.1016/j.pharmthera.2019.107453.

36 Rogers GL, Martino AT, Aslanidi G V., Jayandharan GR, Srivastava A, Herzog RW. Innate immune responses to AAV vectors. Front Microbiol 2011. https://doi.org/10.3389/fmicb.2011.00194.

37 Masat E, Pavani G, Mingozzi F. Humoral immunity to AAV vectors in gene therapy: Challenges and potential solutions. Discov Med 2013.

38 Sharma A, Madhunapantula S V., Robertson GP. Toxicological considerations when creating nanoparticle-based drugs and drug delivery systems. Expert Opin Drug Metab Toxicol 2012. https://doi.org/10.1517/17425255.2012.637916.

39 Scioli Montoto S, Muraca G, Ruiz ME. Solid Lipid Nanoparticles for Drug Delivery: Pharmacological and Biopharmaceutical Aspects. Front Mol Biosci 2020. https://doi.org/10.3389/fmolb.2020.587997.

40 Ndeupen S, Qin Z, Jacobsen S, Estanbouli H, Bouteau A, Igyártó BZ. The mRNA-LNP platform’s lipid nanoparticle component used in preclinical vaccine studies is highly inflammatory. BioRxiv 2021:2021.03.04.430128. https://doi.org/10.1101/2021.03.04.430128.

41 Damase TR, Sukhovershin R, Boada C, Taraballi F, Pettigrew RI, Cooke JP. The Limitless Future of RNA Therapeutics. Front Bioeng Biotechnol 2021. https://doi.org/10.3389/fbioe.2021.628137.

42 Kulkarni JA, Witzigmann D, Thomson SB, Chen S, Leavitt BR, Cullis PR, et al. The current landscape of nucleic acid therapeutics. Nat Nanotechnol 2021. https://doi.org/10.1038/s41565-021-00898-0.

43 Wu SY, Lopez-Berestein G, Calin GA, Sood AK. RNAi therapies: Drugging the undruggable. Sci Transl Med 2014. https://doi.org/10.1126/scitranslmed.3008362.

44 Weng Y, Xiao H, Zhang J, Liang XJ, Huang Y. RNAi therapeutic and its innovative biotechnological evolution. Biotechnol Adv 2019. https://doi.org/10.1016/j.biotechadv.2019.04.012.

45 Aigner A. Perspectives, issues and solutions in RNAi therapy: The expected and the less expected. Nanomedicine 2019. https://doi.org/10.2217/nnm-2019-0321.

46 Wang P, Zhou Y, Richards AM. Effective tools for RNA-derived therapeutics: siRNA interference or miRNA mimicry. Theranostics 2021. https://doi.org/10.7150/thno.62642.

47 Wilson RC, Doudna JA. Molecular mechanisms of RNA interference. Annu Rev Biophys 2013. https://doi.org/10.1146/annurev-biophys-083012-130404.

48 Neumeier J, Meister G. siRNA Specificity: RNAi Mechanisms and Strategies to Reduce Off-Target Effects. Front Plant Sci 2021. https://doi.org/10.3389/fpls.2020.526455.

49 D’Cruz O, Hwang L, Fong A, Ng K, Nam D, Wang W, et al. Preclinical and Clinical Studies on Safety of CEQ508 Bacteria Engineered to Deliver Short-Hairpin RNA to Mediate RNA Interference Against β-catenin in the GI Tract of Patients With Familial Adenomatous Polyposis. Am J Gastroenterol 2017. https://doi.org/10.14309/00000434-201710001-00297.

50 Sautto GA, Kirchenbaum GA, Ross TM. Towards a universal influenza vaccine: different approaches for one goal. Virol J 2018;15:17. https://doi.org/10.1186/s12985-017-0918-y.

51 Monto AS, Malosh RE, Petrie JG, Martin ET. The Doctrine of Original Antigenic Sin: Separating Good from Evil. J Infect Dis 2017. https://doi.org/10.1093/infdis/jix173.

52 Lee LY-H, Ha DLA, Simmons C, de Jong MD, Chau NVV, Schumacher R, et al. Memory T cells established by seasonal human influenza A infection cross-react with avian influenza A (H5N1) in healthy individuals. J Clin Invest 2008. https://doi.org/10.1172/JCI32460.

53 Palese P. Making Better Influenza Virus Vaccines? Emerg Infect Dis 2006;12:http://www.nc.cdc.gov/eid/article/12/1/05-1043_arti. https://doi.org/10.3201/eid1201.051043.

54 Bartlett DW, Davis ME. Insights into the kinetics of siRNA-mediated gene silencing from live-cell and live-animal bioluminescent imaging. Nucleic Acids Res 2006. https://doi.org/10.1093/nar/gkj439.

55 Nel A, Xia T, Mädler L, Li N. Toxic potential of materials at the nanolevel. Science (80-) 2006. https://doi.org/10.1126/science.1114397.

56 Zhang YN, Poon W, Tavares AJ, McGilvray ID, Chan WCW. Nanoparticle–liver interactions: Cellular uptake and hepatobiliary elimination. J Control Release 2016. https://doi.org/10.1016/j.jconrel.2016.01.020.

57 Love SA, Maurer-Jones MA, Thompson JW, Lin YS, Haynes CL. Assessing nanoparticle toxicity. Annu Rev Anal Chem 2012. https://doi.org/10.1146/annurev-anchem-062011-143134.

58 Chvatchko Y, Hoogewerf AJ, Meyer A, Alouani S, Juillard P, Buser R, et al. A key role for CC chemokine receptor 4 in lipopolysaccharide-induced endotoxic shock. J Exp Med 2000. https://doi.org/10.1084/jem.191.10.1755.

59 Ness TL, Ewing JL, Hogaboam CM, Kunkel SL. CCR4 Is a Key Modulator of Innate Immune Responses. J Immunol 2006. https://doi.org/10.4049/jimmunol.177.11.7531.

60 Chen Z, Xie X, Jiang N, Li J, Shen L, Zhang Y. CCR5 signaling promotes lipopolysaccharide-induced macrophage recruitment and alveolar developmental arrest. Cell Death Dis 2021. https://doi.org/10.1038/s41419-021-03464-7.

61 Shi L, Song L, Maurer K, Dou Y, Patel VR, Su C, et al. IL-1 Transcriptional Responses to Lipopolysaccharides Are Regulated by a Complex of RNA Binding Proteins. J Immunol 2020. https://doi.org/10.4049/jimmunol.1900650.

62 Nøhr MK, Kroager TP, Sanggaard KW, Knudsen AD, Stensballe A, Enghild JJ, et al. SILAC-MS based characterization of LPS and resveratrol induced changes in adipocyte proteomics - Resveratrol as ameliorating factor on LPS induced changes. PLoS One 2016. https://doi.org/10.1371/journal.pone.0159747.

63 Nigo YI, Yamashita M, Hirahara K, Shinnakasu R, Inami M, Kimura M, et al. Regulation of allergic airway inflammation through Toll-like receptor 4-mediated modification of mast cell function. Proc Natl Acad Sci U S A 2006. https://doi.org/10.1073/pnas.0510685103.

64 Duan S, Wang N, Huang L, Shao H, Zhang P, Wang W, et al. NLRP3 inflammasome activation involved in LPS and coal tar pitch extract-induced malignant transformation of human bronchial epithelial cells. Environ Toxicol 2019. https://doi.org/10.1002/tox.22725.

65 Liu D, He L, Ding N, Sun W, Qiu L, Xu L, et al. Bronchial epithelial cells of young and old mice directly regulate the differentiation of Th2 and Th17. Biosci Rep 2019. https://doi.org/10.1042/BSR20181948.

66 Boman HG, Monner DA. Characterization of lipopolysaccharides from Escherichia coli K 12 mutants. J Bacteriol 1975. https://doi.org/10.1128/jb.121.2.455-464.1975.

67 Needham BD, Carroll SM, Giles DK, Georgiou G, Whiteley M, Trent MS. Modulating the innate immune response by combinatorial engineering of endotoxin. Proc Natl Acad Sci U S A 2013. https://doi.org/10.1073/pnas.1218080110.

68 Park BS, Song DH, Kim HM, Choi BS, Lee H, Lee JO. The structural basis of lipopolysaccharide recognition by the TLR4-MD-2 complex. Nature 2009. https://doi.org/10.1038/nature07830.

69 Haegens A, Vernooy JHJ, Heeringa P, Mossman BT, Wouters EFM. Myeloperoxidase modulates lung epithelial responses to pro-inflammatory agents. Eur Respir J 2008. https://doi.org/10.1183/09031936.00029307.

70 Brown KC, Montgomery TA. Transgenerational Inheritance: Perpetuating RNAi. Curr Biol 2017. https://doi.org/10.1016/j.cub.2017.03.061.

71 Chang GC, Hsu SL, Tsai JR, Liang FP, Lin SY, Sheu GT, et al. Molecular mechanisms of ZD1839-induced G1-cell cycle arrest and apoptosis in human lung adenocarcinoma A549 cells. Biochem Pharmacol 2004. https://doi.org/10.1016/j.bcp.2004.06.006.

72 Giard DJ, Aaronson S, Todaro G, Arnstein P, Kersey J, Dosik H, et al. In vitro cultivation of human tumors: Establishment of cell lines derived from a series of solid tumors. J Natl Cancer Inst 1973;51:. https://doi.org/10.1093/jnci/51.5.1417.

73 Xiang S, Keates AC, Fruehauf J, Yang Y, Guo H, Nguyen T, et al. In vitro and in vivo gene silencing by TransKingdom RNAi (tkRNAi). Methods Mol Biol 2009. https://doi.org/10.1007/978-1-60327-547-7_7.

74 Keates AC, Fruehauf J, Xiang S, Li CJ. TransKingdom RNA interference: A bacterial approach to challenges in RNAi therapy and delivery. Biotechnol Genet Eng Rev 2008. https://doi.org/10.5661/bger-25-113.

75 Kis Z, Kontoravdi C, Shattock R, Shah N. Resources, production scales and time required for producing RNA vaccines for the global pandemic demand. Vaccines 2021. https://doi.org/10.3390/vaccines9010003.

76 Rosa SS, Prazeres DMF, Azevedo AM, Marques MPC. mRNA vaccines manufacturing: Challenges and bottlenecks. Vaccine 2021. https://doi.org/10.1016/j.vaccine.2021.03.038.

77 Zhu L, Perche F, Wang T, Torchilin VP. Matrix metalloproteinase 2-sensitive multifunctional polymeric micelles for tumor-specific co-delivery of siRNA and hydrophobic drugs. Biomaterials 2014. https://doi.org/10.1016/j.biomaterials.2014.01.060.

78 Bouvier NM, Lowen AC. Animal models for influenza virus pathogenesis and transmission. Viruses 2010. https://doi.org/10.3390/v20801530.

79 Radigan KA, Misharin A V., Chi M, Budinger GRS. Modeling human influenza infection in the laboratory. Infect Drug Resist 2015. https://doi.org/10.2147/IDR.S58551.

80 Bowen LE, Rivers K, Trombley JE, Bohannon JK, Li SX, Boydston JA, et al. Development of a murine nose-only inhalation model of influenza: comparison of disease caused by instilled and inhaled A/PR/8/34. Front Cell Infect Microbiol 2012. https://doi.org/10.3389/fcimb.2012.00074.

